# × *Pseudorhiza nieschalkii* (Senghas) P.F.Hunt nothosubsp. *siculorum* H.Kertész & N.Anghelescu, 2020 AN INTERGENERIC ORCHID HYBRID NEW TO SCIENCE FOUND IN TERRA SICULORUM, ROMANIA

**DOI:** 10.1101/2020.10.21.345462

**Authors:** Nora Eugenia D. G. Anghelescu, Hajnalka Kertész, Nicoleta Constantin, Alexandra Simon-Gruiţa, Georgiana Duţă Cornescu, Maria D. Pojoga, Mihaela I. Georgescu, Sorina A. Petra, Florin Toma

**Affiliations:** University of Agronomic Sciences and Veterinary Medicine of Bucharest, 59 Mărăşti Blvd, District 1, Bucharest, Romania; Bethlen Gábor Middle School & Román Viktor Middle School, Odorheiu Secuiesc, Harghita County, Romania; University of Bucharest, Faculty of Biology, Department of Genetics, Intr. Portocalelor 1-3, Sector 6, Bucharest, Romania

**Keywords:** Pseudorhiza, siculorum, hybrid, orchid, Pseudorchis, Dactylorhiza, hybridization, morphometry

## Abstract

We describe the first reported intergeneric hybrid to naturally occur between two subspecies belonging to different genera, *Dactylorhiza fuchsii* subsp. *sooana* (genus *Dactylorhiza*) and *Pseudorchis albida* subsp. *tricuspis* (genus *Pseudorchis*), as *× Pseudorhiza nieschalkii* (Senghas) P.F.Hunt nothosubsp. *siculorum* H.Kertész & N.Anghelescu, 2020. The hybrid was found and digitally photographed for the first time by Hajnalka Kertész in June, 2020, within Terra Siculorum, in one of the Natura 2000 protected areas, known as Harghita Mădăraş, ROSCI00090. Following detailed morphometric analysis using 67 characters and molecular karyological analyses, we identified this unique specimen as an intergeneric hybrid, new to science. The hybrid, an F_1_ generation plant, most likely representing a single intergeneric pollination event, is phenotypically intermediate between its parental species in most of the characters scored, but it significantly closely resembles *Pseudorchis albida* subsp. *tricuspis* parent. Since the parental species occurred in near proximity (1-10 meters distance), we suggest that the production of this hybrid required a minimum travel distance of ca 1-10 meters, by the pollinators and frequent exchange of pollen between the parental species was very likely. The parental species and the hybrid, which display a considerable synchronicity in their flowering time, overlap in pollinator community, sharing various species of Hymenopterans and Dipterans, very abundant in the heathland. This Terra Siculorum hybrid is thus best described as a rarely occurring intergeneric hybrid that shows strong *Pseudorchis albida* subsp. *tricuspis* parental dominance in inheritance patterns.

## INTRODUCTION

We document the first reported natural occurrence of an intergeneric hybrid between the rare and controversial subspecies, *Dactylorhiza fuchsii* subsp. *sooana* and the alpine species *Pseudorchis albida* subsp. *tricuspis*, the only species present within the hybrid’s close proximity. This previously undescribed alloploid named as *× Pseudorhiza nieschalkii* (Senghas) P.F.Hunt nothosubsp. *siculorum* H.Kertész & N.Anghelescu was discovered by biologist Hajnalka Kertész, on the 30^th^ of June, 2020, during a field trip. The genetic constitution as revealed by the subsequent hybrid karyotyping confirms that its parents are the diploid *Dactylorhiza fuchsii* subsp. *sooana* (genus *Dactylorhiza)* and *Pseudorchis albida* subsp. *tricuspis* (genus *Pseudorchis),* the diploid alloploid genome of the hybrid counting 2n=40 chromosomes.

The hybrid was found in one of the most fascinating heathlands in Tierra Siculorum, one of Natura 2000 protected areas known as Harghita-Mădăraş, very famous for its reach orchid flora. In recent years, our studies encountered several orchid species, usually in substantial numbers such as *Dactylorhiza maculata* (L.) Soó (Heath Spotted-orchid), *Dactylorhiza incarnata* (L.) Soó (Early Marsh-orchid)*, Platanthera bifolia* (L.) Rich. (Lesser Butterfly-orchid), *Dactylorhiza cordigera* subsp. *siculorum* (Soó) Soó (Heart-flowered Marsh-orchid), *Gymnadenia frivaldii* Hampe ex Griseb. (Frivald’s Gymnadenia), *Pseudorchis albida* (L.) Á.Löve & D.Löve (Small White Orchid), *Pseudorchis albida* subsp. *tricuspis* (Beck) E. Klein (Trilobate White Pseudorchis), the dwarf *Dactylorhiza fuchsii* subsp. *psychrophila* (Schltr.) Holub (Cold-loving Dactylorhiza) and rarest and most elusive *Dactylorhiza fuchsii* subsp. *sooana* (Borsos) Borsos (Soó’s Spotted-orchid), all included in our studies. Due to the unusual weather conditions and specific microclimate of the area, which is mainly characterised by cool and rainy summers, most of these orchid species occur in substantial numbers at Harghita-Mădăraş heathland.

The opportunities for this particular hybridisation event to occur are extremely limited, as the parental species *Dactylorhiza fuchsii* subsp. *sooana* and *Pseudorchis albida* subsp. *tricuspis* are rather rare and have contrasting habitat preferences and geographic distributions. To date, Harghita-Mădăraş heathland is the only location where the two subspecies are known to occur in significant, sympatric populations.

This hybrid nothosubspecies belongs to the nothospecies × *Pseudorhiza nieschalkii* (Senghas) P.F.Hunt (hybrid formula: *Dactylorhiza fuchsii* × *Pseudorchis albida*), which is a member of the subtribe Orchidinae Dressler & Dodson, 1960 / Verm., 1955, tribe Orchideae Dressler & Dodson, 1960 / Verm. 1977, subfamily Orchidoideae Lindl., 1826, family Orchidaceae Juss., 1789. The orchid hybrid, × *Pseudorhiza nieschalkii*, was first described in 1968, by German botanist and orchidologist Karlheinz Senghas (1928-2004) and named × *Dactyleucorchis nieschalkii* Senghas [original publication details: *Jahresber. Naturwiss. Vereins Wuppertal* **21-22**: 111 (1968)]. In 1971, British botanist Peter Francis Hunt (born 1936) gave its current name, × *Pseudorhiza nieschalkii* (Senghas) P.F.Hunt [original publication details: *Orchid Rev.* **79** (935): 142 (1971)]. *× Pseudorhiza nieschalkii* nothosubsp. *siculorum* is the first intergeneric hybrid between *Dactylorhiza* and *Pseudorchis* genera, ever mentioned in Romania and the first intergeneric hybrid between *Dactylorhiza fuchsii* subsp. *sooana* and *Pseudorchis albida* subsp. *tricuspis*, ever mentioned in literature. Consequently, we strongly propose this hybrid as new to science and consequently, as a new addition to the Romanian flora.

## MATERIALS AND METHODS

### Location description

The natural area Harghita - Mădăraş is located in the central-eastern part of Harghita County, on the administrative territory of Vlăhiţa and those of the communes Căpâlniţa, Cârţa, Dăneşti, Mădăraş, Racu, Siculeni, Suseni and Zetea (Habitats Directive, 2014). The area was declared a site of Community importance by Order of the Ministry of Environment and Sustainable Development No. 1964 of 13 December 2007 (on the establishment of the protected natural area regime of Sites of Community Importance, as an integral part of the European ecological network Natura 2000 in Romania) and covers an area of 13,373 hectares (List of SCI sites, 2014). Harghita - Mădăraş site is thus designated to protect its biodiversity and preserve the wild flora and fauna within its territory, as well as other natural habitats of community interest located in the protected area. It is located at an altitude between 1,500 and 1,800 m (Mikfalvi & Vifkori, 1979; Marcu, Racz & Cioacă, 1986; Cavruc, 2000).

The site is a natural area covered by deciduous, coniferous and mixed forests, natural meadows, heathlands, bogs (peat bogs) and steppes framed within the alpine bioregion of the Harghita Mountains range. It contains a rich hydrographic network which consist of several lakes and watercourses (Pisota & Bugă, 1976; Ielencz, 2005).

The vegetation of this wetland reserve was dominated by characteristic herbaceous swampy species. *Sphagnum* L. moss was generally abundant, along with various ericaceous shrubs such as *Abies alba* Mill. (Fir), *Picea abies* (L.) H.Karst. (Spruce), *Taxus baccata* L. (Yew), *Fagus sylvatica* L. (Beech), *Quercus petraea* (Matt.) Liebl. (Oak), *Sorbus aucuparia* L. (Mountain Bark), *Fraxinus excelsior* L. (Ash), *Pinus mugo* Turra (Juniper), *Juniperus communis* L. (Juniper), *Sambucus nigra* L. (Sock), *Corylus avellana* L. (Hazel), *Rubus idaeus* L. (Raspberry), *Rosa canina* L. (Rosehip), *Rubus vestitus* Weihe (Blackberry), *Vaccinum myrtillus* L. (Blueberry), *Epilobium nutans* F.W.Schmidt (Willow Herb), *Veronica fruticans* Jacq. (Veronica), *Campanula patula* subsp. *abietina* (Griseb. & Schenk) Simonk. (Bells), to name only a few (Ciocârlan, 2000; Sârbu, Ştefan & Oprea, 2013; EEA, 2014).

This unique hybrid, *× Pseudorhiza nieschalkii* nothosubsp. *siculorum*, was located in a full sun, grassy area, adjacent to a forest margin that covered approximately 1.2 square kilometres. It consisted mainly of a wetland reserve, formed on nutrient poor, acidic to neutral peat substrate. The surface of the soil was moist but not water logged (marshy meadow), mainly covered in *Sphagnum* L. moss.

*D. fuchsii* subsp. *sooana* occurred in moderate numbers and it was found to grow immediately adjacent to the hybrid. Its numbers probably encountered 15-20 plants. The hybrid and *D. fuchsii* subsp. *sooana* parental population were growing within the marshy area of the swamp. The distances between the hybrid and *D. fuchsii* subsp. *sooana* plants were relatively short, the nearest plants occurring within 1-2 meters, while others occurring within 10-15 meters from the hybrid.

*P. albida* subsp. *tricuspis* plants occurred in slightly higher numbers, within the drier parts of the swamp. The distances were somewhat longer, from 10 metres up to 20-40 metres or more, if we take in consideration the distance to the edges of the swamp, where scarce *P. albida* subsp. *tricuspis* plants were found.

We recommend it is imperious to put the entire studied area under strict protection since, shortly after the field studies were performed, the entire heath was completely destroyed by uncontrolled grazing and trampled by the hooves of hundreds of cattle that were walked over the heath.

### Parental species description

#### *Dactylorhiza fuchsii* (Druce) Soó subsp. *sooana* (Borsos) Borsos, 1961

Despite the fact that most of the area was carefully studied, we could not find any typical *D. fuchsii* (Druce) Soó (Common Spotted Orchid) plants, generally described as the tall, vigorous individuals, with the specific deep purple-coloured, highly floriferous inflorescences. Instead, in the close proximity of the hybrid, we found one of its rarest and most elusive subspecies, *D. fuchsii* subsp. *sooana*.

##### Description

The habitus of the plants under study, fits perfectly the original description made in 1959, by the Hungarian botanist Olga Borsos (born 1926), where she described *D. fuchsii* subsp. *sooana* as endemic to the low mountain ranges of Hungary (orig. publ.: *Acta Bot. Acad. Sci. Hung*. **5**: 324, 1959). The Romanian plants were also compared to the typus/holotype registered in the *Flora Hungarica Herbarium* (Reg. No. 3328:152, Rev.: O. Borsos), at the Hungarian Natural History Museum, Bot. Dep. Budapest (data not shown).

*D. fuchsii* subsp. *sooana* is quite similar in habitus (physical aspect) to the type species, *D. fuchsii* (Druce) Soó, although it is slenderer and spindlier, with a slim, solid stem, 7-2 mm in diameter. The plants are in average shorter, reaching up to 30-40 centimetres in height, very rarely 50-60 centimetres (*D. fuchsii* plants are very sturdy, in some cases reaching up to 90-100 centimetres). The lowermost leaf (basal leaf) is elongated with an oval, roundish tip. It is distinguished from *D. maculata* (L.) Soó by the oval tip of the lowest leaf, the leaves being broadest in the upper half. The 5-8 cauline leaves are lanceolate to narrow-lanceolate, elongate, deep green, always brownish-purple spotted on the upper side (sometimes the spots are oval to flat-elongated in shape), wider around the middle/upper half. The inflorescence is elongated, lax to dense and floriferous and may bear up to 35-40 flowers. The flowers are medium-sized, pure white. In some rare cases, the lateral sepals and the spur are faintly tinged pink. The sepals are spreading obliquely to subtend angle of c.45° relative to the hood. The lateral petals and the dorsal sepal form a loose hood over the fertile anther (gynostemium). The labellum is white, three-lobed and spreading. The median lobe is prominent, triangular almost as broad as and longer than the lateral lobes, sometimes with a rounded tip. The markings on the labellum consist exclusively of few, short, irregular, purple-reddish dots, streaks, lines or irregular loops. One of the most important characteristics of this subspecies is that the purple markings never form continuous loops or circles. The lateral lobes are scalloped, flat, with unmarked lateral margins. The spur is slim, straight to horizontal, slightly curved downwards at tip. The spur secrets no nectar. The flower bracts are shorter than flowers, more or less equal to the ovary, except at the base of the inflorescence, where they are slightly longer than the ovaries, but shorter than the flowers.

##### Chromosome number

2n=40 [similar to type species *D. fuchsii* (Druce) Soó]

##### Habitat

*D. fuchsii* subsp. *sooana* prefers swampy areas, very wet meadows, humid forest margins, on alkaline to slightly acidic substrates (in present study, it occurred on moist substrate of *Sphagnum* L. moss), up to 1,700 meters altitude.

##### Distribution

In Romania, *D. fuchsii* subsp. *sooana* occurs exclusively within the protected area Harghita-Mădăraş, in medium homogenous populations (Kreutz, 2014). Beside Hungary and Romania, *D. fuchsii* subsp. *sooana* was also mentioned to occur in the White Carpathians of Ukraine (Loya, 2015), Czech Republic and Slovakia (Danihelka *et al.*, 2012)

##### Etymology

The *generic name*, *Dactylorhiza*, is a compound term that originates in the ancient Greek words *dáktylos* (finger) and *rhíza* (root), *ad litteram* meaning *finger-shaped* or *finger-like roots*, a reference to the *palmate*, *two-* to *five-lobed tubers* that resemble the *fingers of a human palm*, a characteristic of all the species belonging to this genus.

The *specific epithet*, *fuchsii*, was given in honour of Leonhart/Leonhard Fuchs (1501-1566), a German physician and botanist, hence its common name, Fuchs’ Dactylorhiza. Taxa with the specific epithet, *fuchsii*, commemorate his name. Due to the strongly *maculated leaves*, this species is also known by its vernacular names, the Common Spotted Orchid, the Spotted Orchid or the Marsh Spotted Orchid.

The *infraspecific epithet* (*subspecies epithet*), *sooana*, was given in honour of Károly Rezső Soó von Bere (1903-1980), a Hungarian botanist, professor at the University of Budapest, born in Odorheiu Secuiesc, Romania, hence its vernacular name, Soó’s Spotted-orchid. Taxa with the specific epithet, *sooana*, commemorate his name.

#### *Pseudorchis albida* (L.) Á.Löve & D.Löve subsp. *tricuspis* (Beck) E.Klein, 2000

##### Description

*P. albida* subsp. *tricuspis* is a slender, tall plant that may reach 20-35 (occasionally 40-45) centimetres in height. The stem is erect, greenish, 3.4-3.9millimetres in diameter. The basal leaf is widest around the middle. The 5-8 cauline leaves are shiny-green to yellowish-green, oblong-obovate, widest around the middle, briefly pointed. The cauline leaves (stem leaves) are narrower, lanceolate, the upper ones becoming, pointed, bract-like. The inflorescence is up to 6-9 centimetres long, dense, narrow-cylindrical, very floriferous, bearing up to 35-65 small, yellowish-green flowers. The flowers are small, whitish to yellowish-green with complete or partial resupination (Jersáková *et al.*, 2011). The lateral petals and sepals are very similar and form a tight hood over the anther/gynostemium. The labellum is the main distinctive feature of the species. It is yellow, flat, deeply three-lobed with all three lobes of equal lengths. Spur is 2.5-3.4 millimetres long, cylindrical, slightly curved downwards, yellowish-white, blunt and secretes abundant nectar. The flower bracts of the lower/basal flowers of the inflorescence are narrow-lanceolate, longer than the top ones (some are a significantly longer than the flowers). In general, bracts are longer than the ovaries, but shorter than the flowers (Delforge, 2006).

##### Chromosome number

2n=42 (Klein, 2000; Jersáková *et al.*, 2011).

##### Habitat

In Romania it grows in well-drained to wet meadows, grasslands, pastures, forest edges and rocky slopes, alpine pastures and meadows, at 700-2,300 meters altitude. It prefers a wide range of soil conditions from the acidic moist substrata of *Sphagnum* bogs to alkaline/calcicolous (Klein, 2000), well drained soils on carboniferous limestone (Summerhayes, 1951). Plants having attributes of *P. albida* subsp. *tricuspis* grow on acidic soils in the Czech Republic (Jeřabková *et al.*, 2006).

##### Distribution

*P. albida* subsp. *tricuspis* is a palearctic species, covering boreal alpine, subalpine and temperate zones, from Europe to the Russian Far East, to the Northern Urals and Kamchatka and from Eastern Canada to Greenland, but not in Siberia.

##### Etymology

The Catholic priest and renowned Italian botanist Pier Antonio Micheli (1679-1737) was the first to illustrate and use the name *Pseud-orchis* (*false orchid*) for this genus, in his work *Nova Plantarum Genera* (1729), in order to indicate the similar appearance of the plant to the members of the genus *Orchis*.

The generic name, *Pseudorchis*, originates in the ancient Greek words *pseûdos* (false) and *órkhis* (*testicle* or *a plant with roots like testicles*), a word used for the first time by Theophrastus (372-286 B.C.E.), in his book *Historia Plantarum (Enquiry into Plants)*. Consequently, *Pseudorchis* may be interpreted as the *false-orchid*, which refers to the shape of its *root-tubers* that are not *round*, like all the other *Orchis* species, but *bifid* or *palmate*, hence the vernacular name of these species, the False Orchid.

The specific epithet, *albida*, has its origin in the Old Latin word *alba*, which means white and refers to the *whitish flowers* of this subspecies, hence its vernacular names, the Small White Orchid, the White Mountain Orchid or the Bright White Pseudorchis.

The subspecific (infraspecific) epithet, *tricuspis*, originates in the ancient Greek words *tría* (three) and *cuspis* (tip) meaning *with three tips*, a reference to the three prominent lobes of the labellum, hence its potential vernacular names, the Small White Trilobate Orchid or the Trilobate White Pseudorchis.

### Flowering times

The flowering times of parental species and the resulted hybrid overlapped entirely, from mid-June to late July. At the time when the studies were performed, all three species were at the peak of anthesis (flowering times). In some parts of the swamp, some *D. fuchsii* subsp. *sooana* individuals were slightly off the peak of anthesis, which indicates that *D. fuchsii* subsp. *sooana* parent might have an earlier blooming time, with approximately 3-5 days before *P. albida* subsp. *tricuspis* parent. Nevertheless, this slight synchronicity mismatch is insignificant and does not influence the frequency of cross-pollination events, which might potentially result in the occurrence of the hybrid.

#### General descriptions of the hybrid *× Pseudorhiza nieschalkii* (Senghas) P.F.Hunt nothosubsp. *siculorum* H.Kertész & N.Anghelescu and parental species comparison diagrams

In general, primary hybrids (F_1_ generation) appear phenotypically intermediate between the parental species. Nevertheless, in the case of *× Pseudorhiza nieschalkii* nothosubsp. *siculorum*, a rather more notable influence of *P. albida* subsp. *tricuspis* parent is evident.

### Morphometric/biometric methods

Given that the parental species differ considerably in morphology, the identification of any hybrids between them, appear relatively straight-forward (the images are very explicit). In most cases, the phenotypical characters/traits of the hybrid appear to resemble more *P. albida* subsp. *tricuspis* parent.

It should be emphasised that a positive determination of hybrids implies a deep knowledge of the variation of the parental species and a deep characterisation of the biotope in which they are found. Therefore, for the correct identification of a hybrid, it is imperious that at least one unequivocal character of each parental partner is demonstrable and cannot come from the other supposed partner (Bateman & Hollingsworth, 2004).

In order to describe this unique hybrid as comprehensive as possible, a wide range of characters were taken in consideration and biometrically/morphometrically analysed. All morphological measurements were undertaken in the field. In total, 68 morphological characters were measured directly (Table 1). Morphological characters used for analysis included most of the characters used by Bateman & Denholm (1985), Bateman & Hollingsworth (2004) and by Pedersen & Hedrén (2010). Special attention was given to the characters that proved to be taxonomically informative and those that involve the details of labellum morphology. The quantitative measurements encompass all organs except the root-tubers and the reproductive organs (fused forming the gynostemium). Measurements are examples of the parental plants and of the hybrid.

**Table 1.** Morphometric comparison of the putative parental species and the unique resulted hybrid. The quantitative measurements (in millimetres unless otherwise stated) encompass all organs except the root-tubers and gynostemium.

| Vegetative & Floral Organs | Characters / Features (millimetres) | *Dactylorhiza fuchsii* subsp. *sooana* | × *Pseudorhiza nieschalkii* nothosubsp. *siculorum* | *Pseudorchis albida* subsp. *tricuspis* |
| --- | --- | --- | --- | --- |
| 1. Stem & Inflorescence |  |  |  |  |
|  | Overall height | 333/400 | 285 | 300/350 |
|  | Stem diameter | 3.5/3.7 | 3.2 | 3.4/3.9 |
|  | Stem anthocyanins | moderate/strong | traces apically | absent |
|  | Inflorescence length | 53/60 | 67 (23 % of stem length) | 60/80 (90) |
|  | Inflorescence shape | conical 25/40 | cylindrical, flat topped | elongated, cylindrical, acuminate top |
|  | No. of flowers | 25/40 | 28 | 45/65 |
| 2. Leaves |  |  |  |  |
|  | No. of basal leaves | 1, rounded at tip | 1, widest around the middle | 1, widest around the middle |
|  | Distribution of sheathing leaves on stem | Even/lower half | basal/even | denser basally/even |
|  | Longest leaf posture | Spreading, to subtend angle of c.45° relative to the stem | spreading horizontally to subtend angle of c.90° relative to the stem | Spreading, to subtend angle of c.45° (90° in some cases) relative to the stem |
|  | No. of sheathing leaves | 5/8 | 4 | 5/8 |
|  | No. of non-sheathing leaves | 1 | 1 | - |
|  | Length of longest leaf | 70/74 | 65 | 22/72 |
|  | Width of longest leaf | 12/13.2 | 11 | 11/15 |
|  | Outline shape of longest leaf | linear-lanceolate, broadest in the upper half | linear-lanceolate | linear-lanceolate, widest around the middle |
|  | Leaf conduplicate | strong | faint/absent | strong |
|  | Apex hooding | moderate/faint | lacking pronounced apical hooding | moderate/strong |
|  | Leaf colour | deep green | vivid/shiny green | shiny, light to deep green |
|  | Leaf dorsal side | deep green | green | deep green |
|  | Leaf ventral side | green | green | deep green, veined |
|  | Leaf margins | entire | entire | entire |
|  | Leaf markings | brownish-purple spotted on the upper side | unmarked | unmarked |
|  | Upper leaves | shorter, narrow-lanceolate, acuminate | bract-like | shorter, narrow-lanceolate, acuminate, bract-like |
| 3. Bracts & Ovary |  |  |  |  |
|  | Length of basal bracts | 8/13 exceeding the ovaries | 14/16, exceeding flowers | 10/16, exceeding the flowers |
|  | Width of basal bracts | 1.5/2.3 | 1.9/2.5 | 2.8/3.2 |
|  | Length of floral bracts | 11/8, equal/slightly longer than the ovaries | 13/10, longer/equal to the ovaries | 12/9, longer than the ovaries |
|  | Width of floral bracts | 1.2/2.1 | 1.1/1.4 | 2.1/3 |
|  | Texture of bracts | membranous/robust | robust/membranous | robust |
|  | Bract anthocyanins | strong | traces present in the top bracts | absent |
|  | Marginal wall thickness | thin | moderate | thick |
|  | Ovary length | 5.9/7.3 | 5.2/7 | 4.3/5.7 |
|  | Ovary diameter | 1.2/2.2 | 1.2/1.6 | 2.1/3.2 |
|  | Ovary anthocyanins | strong | traces | absent |
| 4. Sepals & Lateral Petals |  |  |  |  |
|  | Lateral sepals position | spreading to subtend angle of c.45° relative to the hood | spread horizontally, laterally relative to the hood | tightly connivant forming the hood |
|  | Lateral sepals connivant | no | no | yes |
|  | Sepal fusion from base (%) | - | 5/10 | 90/95 |
|  | Sepal apex | acuminated | oval/blunt | oval/roundish |
|  | Sepal iridescent green pigment | absent | absent | present |
|  | Median sepal length | 5.2/6/7 | 4.8/5.7 | 4.3/5.8 |
|  | Median sepal width | 3.2/4.6 | 2.7/3.7 | 2.4/3.8 |
|  | Lateral sepals width | 1.7/2.4 | 2.1/2.9 | 2.5/3.9 |
|  | Lateral sepals length | 4.6/5.7 | 4.7/5.8 | 3.9/4.5 |
|  | Lateral petals width | 2.4/3.2 | 1.7/2.3 | 2.4/3.7 |
|  | Lateral petals length | 3.9/4.8 | 3.8/4.6 | 3.8/4.7 |
| 5. Labellum |  |  |  |  |
|  | Outline shape | orbicular, nearly flat | orbicular, deeply three-lobed, nearly flat | deeply three-lobed |
|  | Lobes | Large, scalloped | three, triangular-elongated, equal in length | three, thin-elongated, equal in length |
|  | Median lobe length | 2.1/2.3 | 2.9/3.1 | 2.3/3.4 |
|  | Lateral lobes length | 1.4/1.9 | 2.6/2.9 | 2.1/3.3 |
|  | Sinuses separating the three lobes | shallow | very deep, c.2.1/2.3 | very deep, c.2.2/2.4 |
|  | Width | 3.9/4.8 | 1.4/2.3 | 0.9/1.2 |
|  | Lateral lobe reflexion | slightly reflexed upwards | moderate to slightly recurved backwards | moderate to slightly recurved backwards |
|  | Central lobe apex | shallow, oval-roundish | acuminate, entire, flat | acuminate-pointy, entire, flat |
|  | Central lobe width | 4.2/5.3 | 1.8/2.1 | 1.6/2 |
|  | Base colour of labellum | white | white with a yellowish splash at spur entrance | yellowish-green |
|  | Centre colour | white, marked with scattered with dots and streaks that never form circular loops | whitish, convex, uniformly dotted | yellowish, unmarked |
|  | Margins colour | white | white | yellowish |
|  | Markings type | purple dots and streaks that never form circular loops | scattered faint pinkish dots & streaks, never forming circular loops | uniformly yellow, unmarked |
|  | Surface markings papillate | moderately | moderately | no |
|  | Markings distribution in centre | Centrally, lateral sides of the lobes white | uniformly scattered | circular |
|  | Markings contrast | strong | weak | - |
|  | Lateral lobe indentation | entire, scalloped | entire/moderate indentation, scalloped, slightly recurved | entire |
| 6. Spur |  |  |  |  |
|  | Spur length | 3.2/4/5 | 3.2/3.8 | 2.5/3.4 |
|  | Spur shape | cylindrical thin, elongated, shorter than the ovary | cylindrical, thin, with a roundish tip, c. ½ ovary length | Cylindrical, blunt, whitish, c. ½ ovary length |
|  | Spur width entrance | 2.2/2.8 | 1.3/2 | 1.1/2 |
|  | Spur width halfway | 1.4/1.7 | 1.2 | 1.1/1.2 |
|  | Spur down-curvature | straight to slightly down-curved | slightly down-curved | slightly down-curved |
| 7. Nectar |  |  |  |  |
|  | Presence | no | - | yes |
|  | Amount | - | - | abundant |
| 8. Smell |  |  |  |  |
|  | Fragrance type | vaguely fruity | - | moderate/strong, sweet, vanilla like |

**Table 2.** Chromosome numbers of the specimens studied.

| Taxon | Distance from hybrid (m) | Chromosome no. |
| --- | --- | --- |
| *D. fuchsii* subsp. *sooana* | 2m | 2n=40 |
| *D. fuchsii* subsp. *sooana* | 5m | 2n=40 |
| *P. albida* subsp. *tricuspis* | 7m | 2n=42 |
| *P. albida* subsp. *tricuspis* | 10m | 2n=42 |
| × *P. nieschalkii* nothosubsp. *siculorum* no. 28-2 | - | 2n=40 (±2) |
| × *P. nieschalkii* nothosubsp. *siculorum* no. 30-1 | - | 2n=40 (±2) |
| × *P. nieschalkii* nothosubsp. *siculorum* no. 31-4. | - | 2n=40 (±2) |

### Chromosome counts

Explants, 3-4 millimetres long adventitious roots tips (meristematic root tips), were sampled from the hybrid and its parents. Chromosome counts were carried out using preparations of dividing cells (at metaphase) from the apical portion of adventitious roots. The root tips were pre-treated in a 0.5% colchicine and incubated for 2 h at 10–15 °C. These were transferred to Clark fixative 3:1 (v/v) (3 parts absolute ethanol and 1part glacial acetic acid) at 10–15 °C for 30 min and then 3–4 °C for 48 h. Further, meristematic root tips were hydrolysed at 60°C in 1 N HCl for 60 min and stained in Feulgen stain. For cytogenetic analysis, the stained root tips were macerated on a microscope slide in a drop of acetocarmine for 1–2 min, covered with a coverslip and squashed manually (classical squash method in acetocarmine).

## Results

### Morphometric comparisons

The morphometric results proved to be partially asymmetric. The most effective diagnostic characters, such as labellum morphology, placed the hybrid closer to the *P. albida* subsp. *tricuspis* parent, rather than *D. fuchsii* subsp. *sooana* parent.

#### Habitus

The hybrid was tall and sturdy, reaching up to 28.5 centimetres, intermediate or even similar in size to the parents, which both usually range between 25-40 centimetres in height (taller parental species that may reach up to 40-60 centimetres are very rare and were not present among the individuals included in this study).

#### Stem

The stem is slender and slim, vividly-green coloured. It presents no significant purple pigmentation at the tip (only trace amounts of anthocyanins), resembling exclusively *P. albida* subsp. *tricuspis* parent.

#### Leaves

The hybrid presents 1 unspotted lanceolate basal leaf and 4 narrow-lanceolate cauline leaves (stem leaves), the upper one being very reduced, bract-like, not reaching the inflorescence. The leaves present no maculae or any other pigmentation, a character inherited exclusively from *P. albida* subsp. *tricuspis* parent that has unspotted, bright-green leaves, therefore, the inheritance of leaf characters proved to be asymmetrical.

#### Inflorescence

The inflorescence is not as floriferous as *P. albida* subsp. *tricuspis* parent. It resembles mostly *D. fuchsii* subsp. *sooana* parent, both in shape - cylindrical, with a flat top, and number of flowers - it is less floriferous, bearing approximately 28 medium sized flowers.

#### Flowers

The flowers’ size of the hybrid are intermediate between the parents and range mostly in the median range of the parental sizes (length *×* width), being slightly larger than those of *P. albida* subsp. *tricuspis* parent and smaller to almost equal in size to *D. fuchsii* subsp. *sooana* parent.

#### Bracts

The flower bracts are slightly variable in size, the lower bracts being longer than the flowers, while the top bracts being slightly longer than the ovaries but shorter than the flowers. Overall the bracts resemble mostly *P. albida* subsp. *tricuspis* parent, since *D. fuchsii* subsp. *sooana* parent has bracts shorter to almost equal to the ovaries. The top flower bracts show a faint purple pigmentation, a trait mildly inherited exclusively from *D. fuchsii* subsp. *sooana* parent and totally absent from *P. albida* subsp. *tricuspis* parent.

#### Sepals & lateral petals

The sizes are again, intermediary between the parental species, but the colour and pinkish markings are mainly inherited from *D. fuchsii* subsp. *sooana* parent. The background colour of the tepals is white with faint purple-pink dots and streaks on the lateral sepals, a characteristic exclusively inherited form *D. fuchsii* subsp. *sooana* parent, with the mention that hybrid’s pigmentation is less pronounced, the anthocyanins being expressed in far smaller amounts.

#### Helmet/hood

The lateral petals and sepals that construct the helmet, which protects and covers the reproductive organs (fused to form the gynostemium) are not so tightly folded. The lateral sepals significantly resemble *D. fuchsii* subsp. *sooana* parent as they are laterally/horizontally spreading, hence the loose helmet is formed mainly by the dorsal sepal and lateral petals, completely different from the tight helmet formed from the highly connivant sepals and lateral petals in *P. albida* subsp. *tricuspis* parent.

#### Labellum

The morphology of the labellum is particularly interesting as the parents differ considerably in the labellum size and shape. The overall shape of the labellum closely resembles *P. albida* subsp. *tricuspis* parent. The labellum is deeply three-lobed with all lobes almost equal in *length and width,* a characteristic exclusively inherited from *P. albida* subsp. *tricuspis* parent, although they are wider than those of *P. albida* subsp. *tricuspis* parent, a feature partly inherited from *D. fuchsii* subsp. *sooana* parent, which has scalloped, rounded labellar lobes.

#### Labellar colour and markings

Labellum colour is intermediate between the parents. The *yellowish* background colour of the labellum base (spur entrance) is inherited from *P. albida* subsp. *tricuspis* parent, which has yellowish-green flowers. The white background colour of the labellum and the *faint pinkish marks* represent the distinctive features of this nothosubspecies, clearly inherited from *D. fuchsii* subsp. *sooana* parent. The pinkish markings are distinctively fainter but resemble *D. fuchsii* subsp. *sooana*’s patterns of scattered dots and streaks that never form circular loops.

#### Spur

The spur mostly resembles *D. fuchsii* subsp. *sooana* parent, being slightly longer and slenderer than in *P. albida* subsp. *tricuspis* parent (but shorter than the ovary), cylindrical and slightly curved downwards.

### Chromosome counts

Stained cells were viewed using Micros Microscope, 100X lenses and photographed with Optika microscopes camera, lenses 0,45X. Images were analysed using OptikalSview, developed by OPTIKA Microscopes Software.

Chromosomes were counted for parental species and hybrid and confirmed expected numbers: *D. fuchsii* subsp. *sooana*: 2n=40, *P. albida* subsp. *tricuspis*: 2n=42 and *× P. nieschalkii* nothosubsp. *siculorum* 2n=40 (±2), which makes the intergeneric hybrid a diploid (alloploid) nothosubspecies. These findings also comply with the fact that the parental species have close values of the chromosome number, 2n=40 & 42, allowing them to cross-pollinate relatively easily.

## Discussion

The intergeneric hybrid plant was found in the western area of Harghita-Mădăraş heath, by Hajnalka Kertész, on 30^th^ of June 2020, at 08:30:19am (Fig. 3). Its likely parentage was immediately recognised, its putative parents growing within 2 and 10 meters respectively, but its greater significance as a hybrid combination new to science was not fully elucidated at that moment. Therefore, digital images were taken by Hajnalka Kertész, but neither detailed measurements nor samples suitable for chromosomal analysis were obtained. Very soon after, digital images were sent to Nora Anghelescu who replied almost immediately, not only confirming that it was indeed a hybrid plant but also pointing out that this hybrid combination was new to science. Following an urgent request by Nora Anghelescu, several trips followed, trips during which the hybrid and the putative parents were analysed, measured and photographed in great detail.

**Fig. 1.**
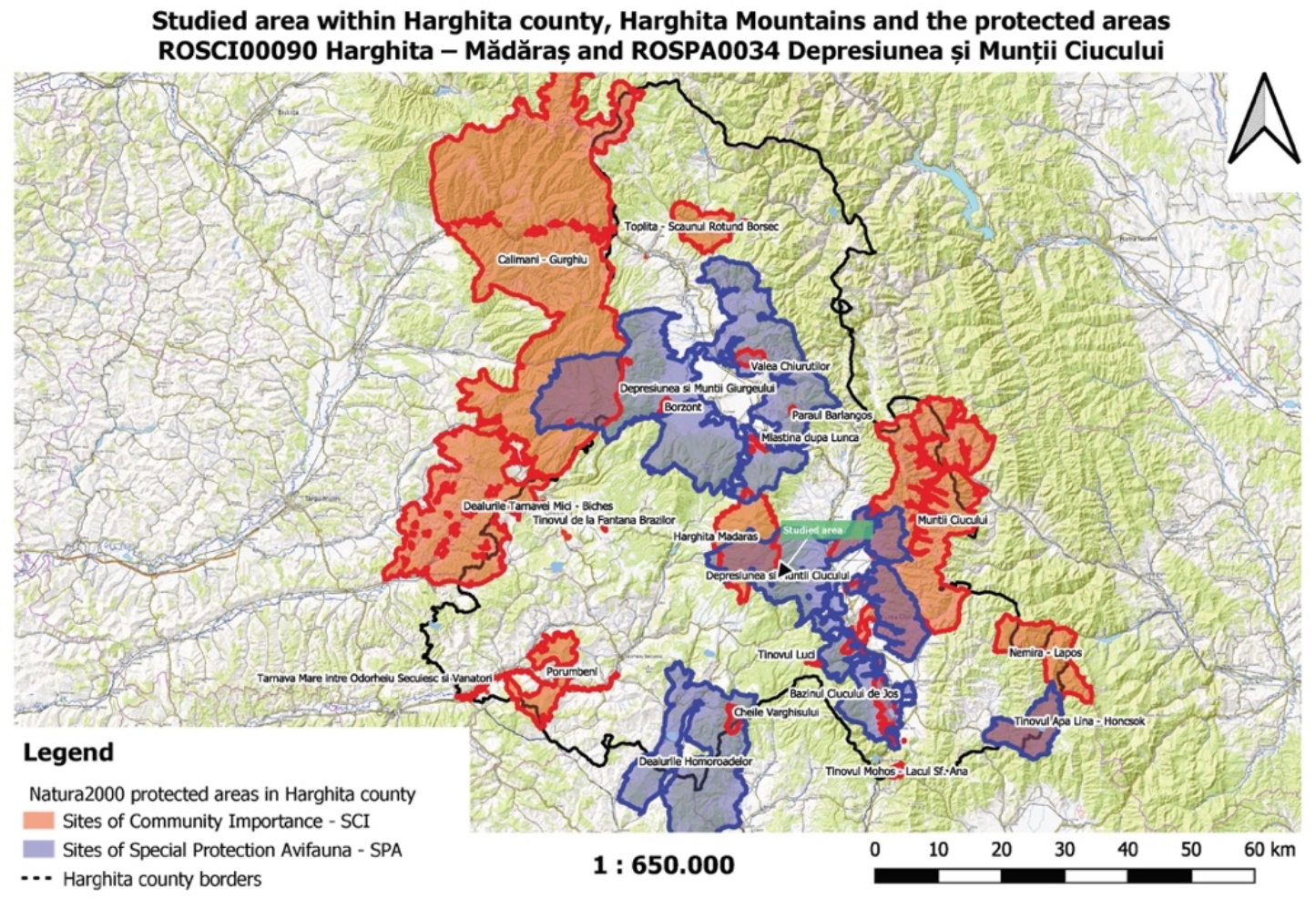
The map of the studied area within Harghita county, Harghita Mountains and the protected areas ROSCI00090 Harghita – Mădăraş and ROSPA0034 Depresiunea şi Munţii Ciucului (map used by permission of National Agency for Protected Areas Harghita Territorial Service)

**Fig. 2.**
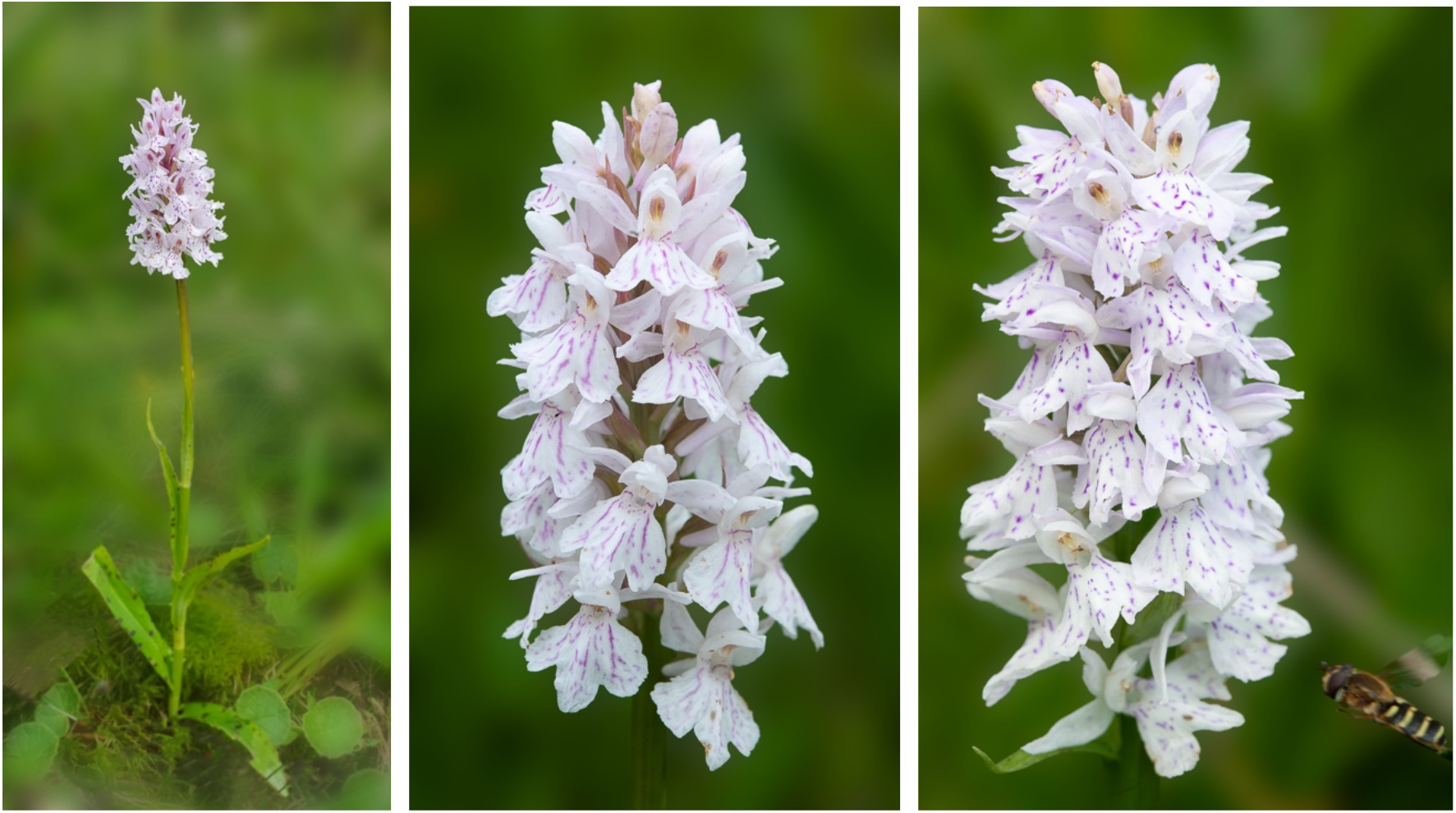
*Dactylorhiza fuchsii* (Druce) Soó subsp. *sooana* (Borsos) Borsos entire plant in its natural habitat (A), details of the inflorescences (B, C) One of the inflorescences are visited by hoverfly of the family Syrphidae Latreille, 1802. The hoverflies, which are true bee-mimics, are very frequent visitors of *D. fuschii* subsp. *sooana*, although it has not been shown to be its real pollinators, as of yet. Photos A-C © N. Anghelescu originals

**Fig. 3.**
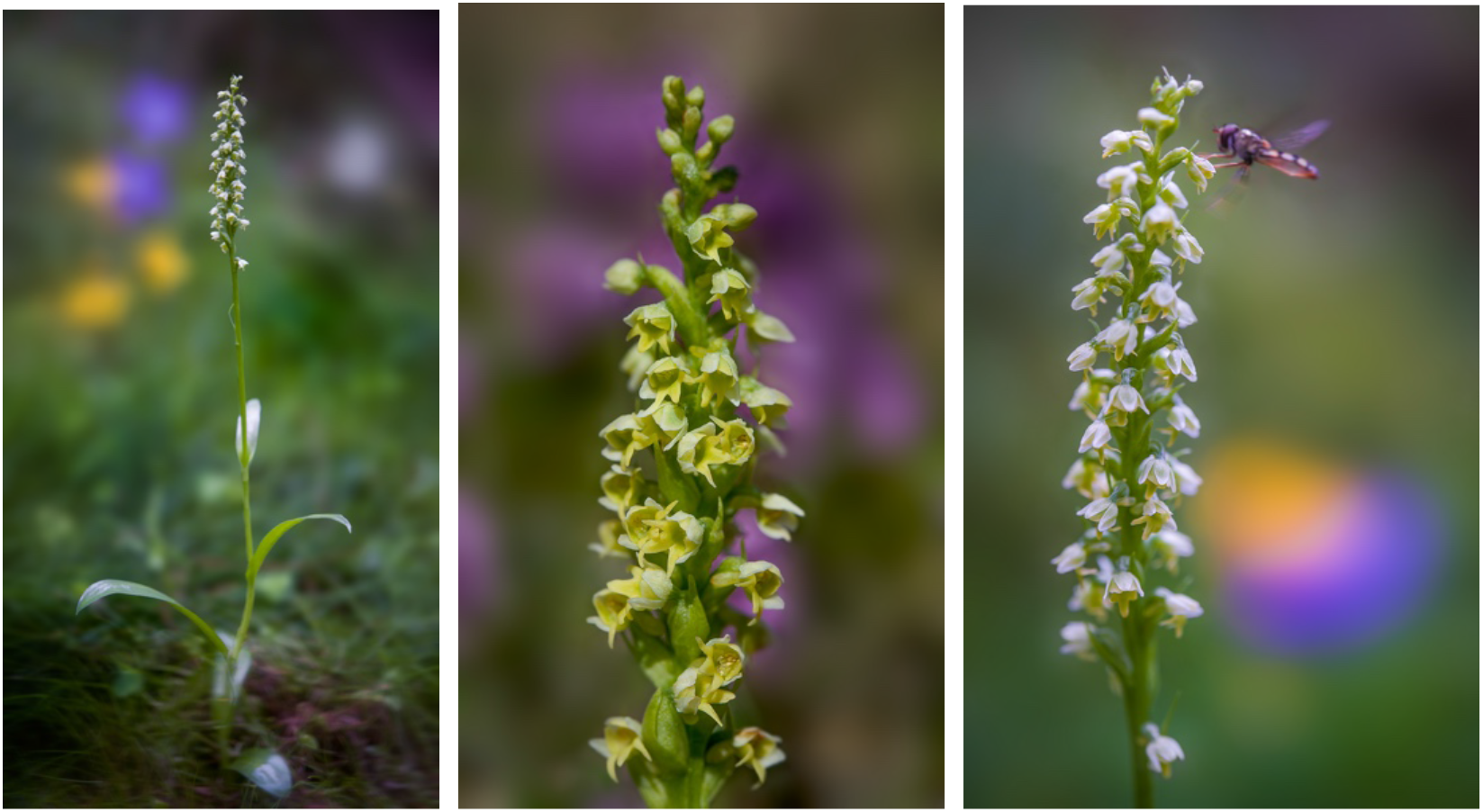
*Pseudorchis albida* (L.) Á.Löve & D.Löve subsp. *tricuspis* (Beck) E.Klein entire plant in its natural habitat (A), details of the inflorescences (B, C) One of the inflorescences are visited by hoverfly of the family Syrphidae Latreille, 1802. The hoverflies, which are true bee-mimics, are also very frequent visitors of *Pseudorchis albida* subsp. *tricuspis*, although it has not been shown to be its real pollinators, as of yet. Photos A-C © N. Anghelescu originals

**Fig. 4.**
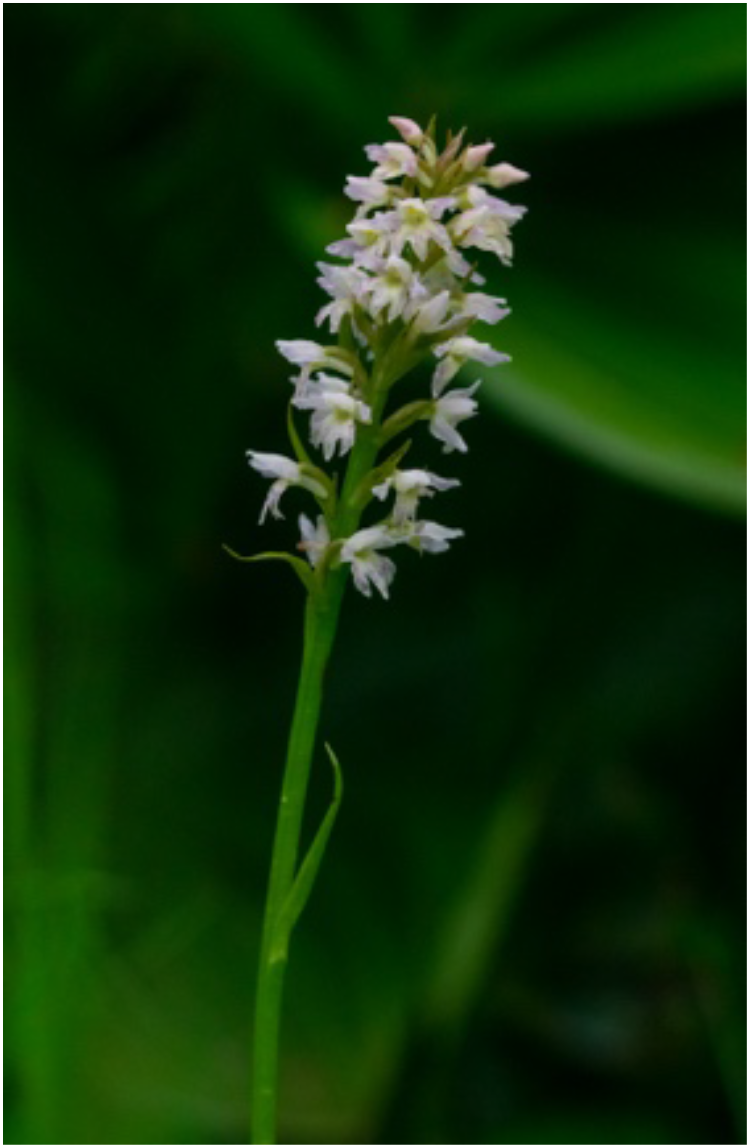
This is the first image of *× Pseudorhiza nieschalkii* nothosubsp. *siculorum* ever taken by H. Kertész at 08:39:19am, on 30^th^ of June, 2020. While researching the existing orchid species of the heathland, she spotted an oddly looking plant, which, from a distance, resembled *P. albida* subsp. *tricuspis.* She approached the plant and, after parting the dense grass that surrounded it, she realised it was something more than a simple *P. albida* subsp. *tricuspis*. Minutes later digital images were sent to N. Anghelescu who confirmed the putative parents and, in that regards, suggested that the hybrid may actually be new to science. Photo © H. Kertész original

**Figure 5.**
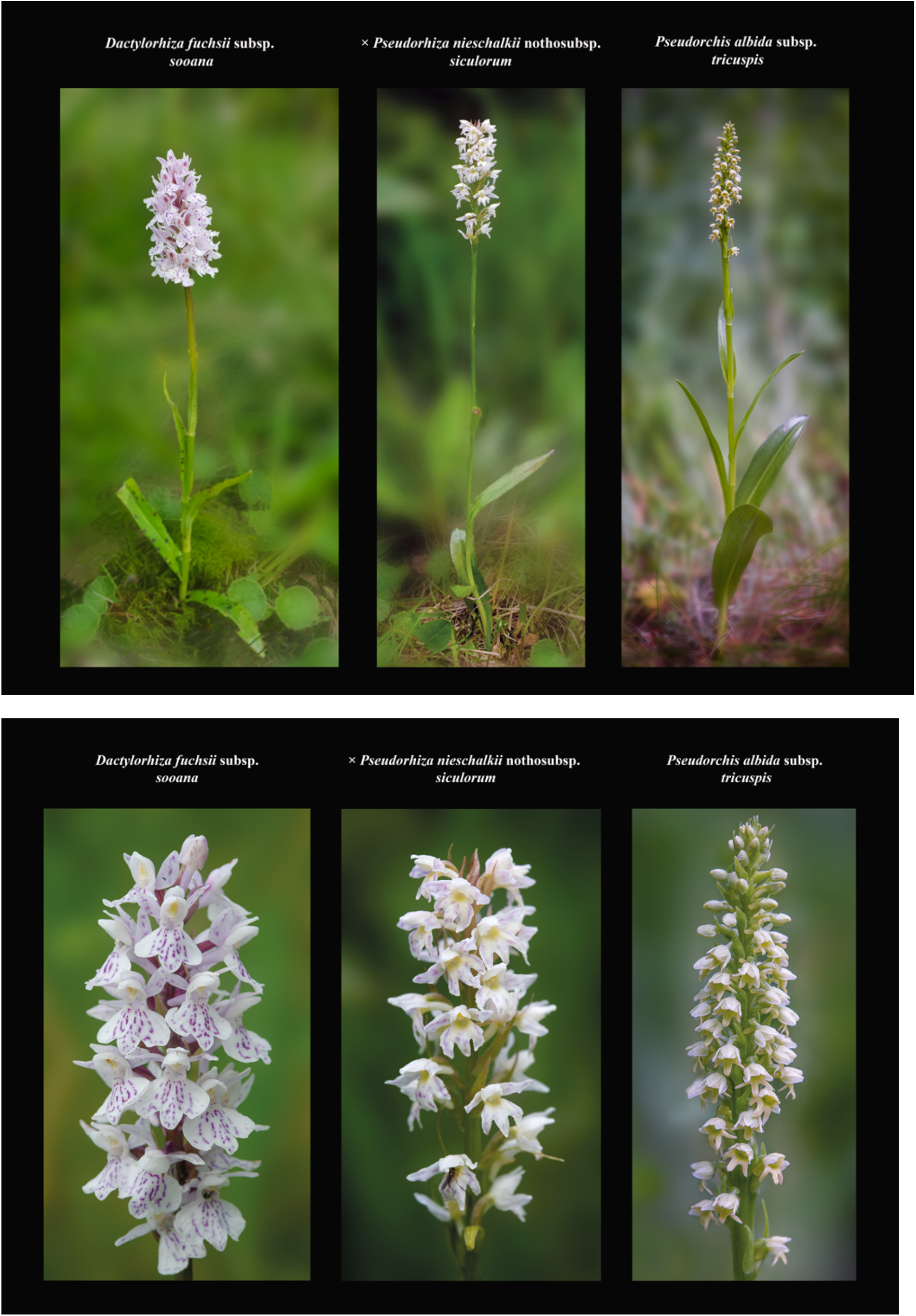
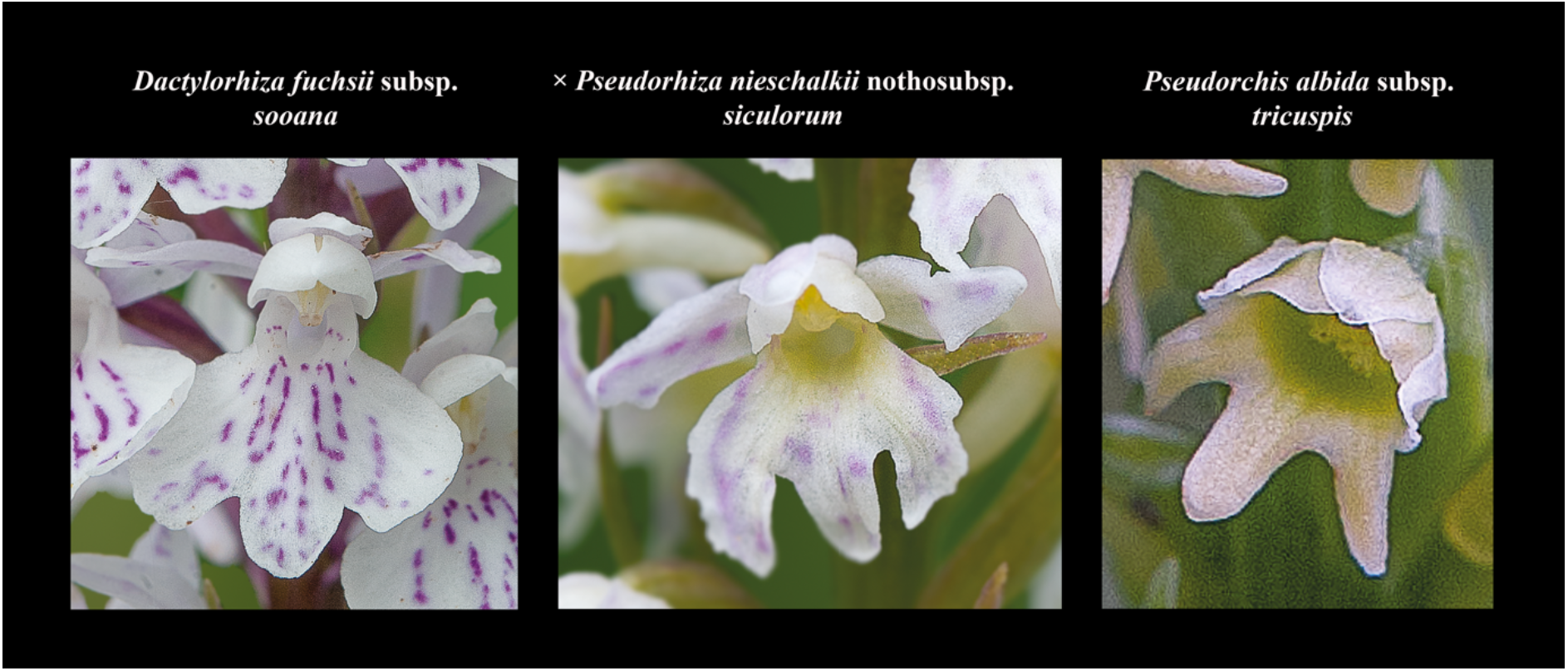
Morphological comparisons of *× Pseudorhiza nieschalkii* nothosubsp. *siculorum* and parents (A-D). The following three comparison diagrams show the intermediate characters of the hybrid’s habitus (A), inflorescence (B) and flowers (C), compared to those of its parents. The hybrids are almost as tall as the parents. The hybrid’s inflorescence is less floriferous and flat topped, resembling more that of *D. fuchsii* subsp. *sooana* parent. The flowers’ labellum is deeply three-lobed, very similar to *P. albida* subsp. *tricuspis*’ labella. Photos A-C © N. Anghelescu originals

**Fig 6.**
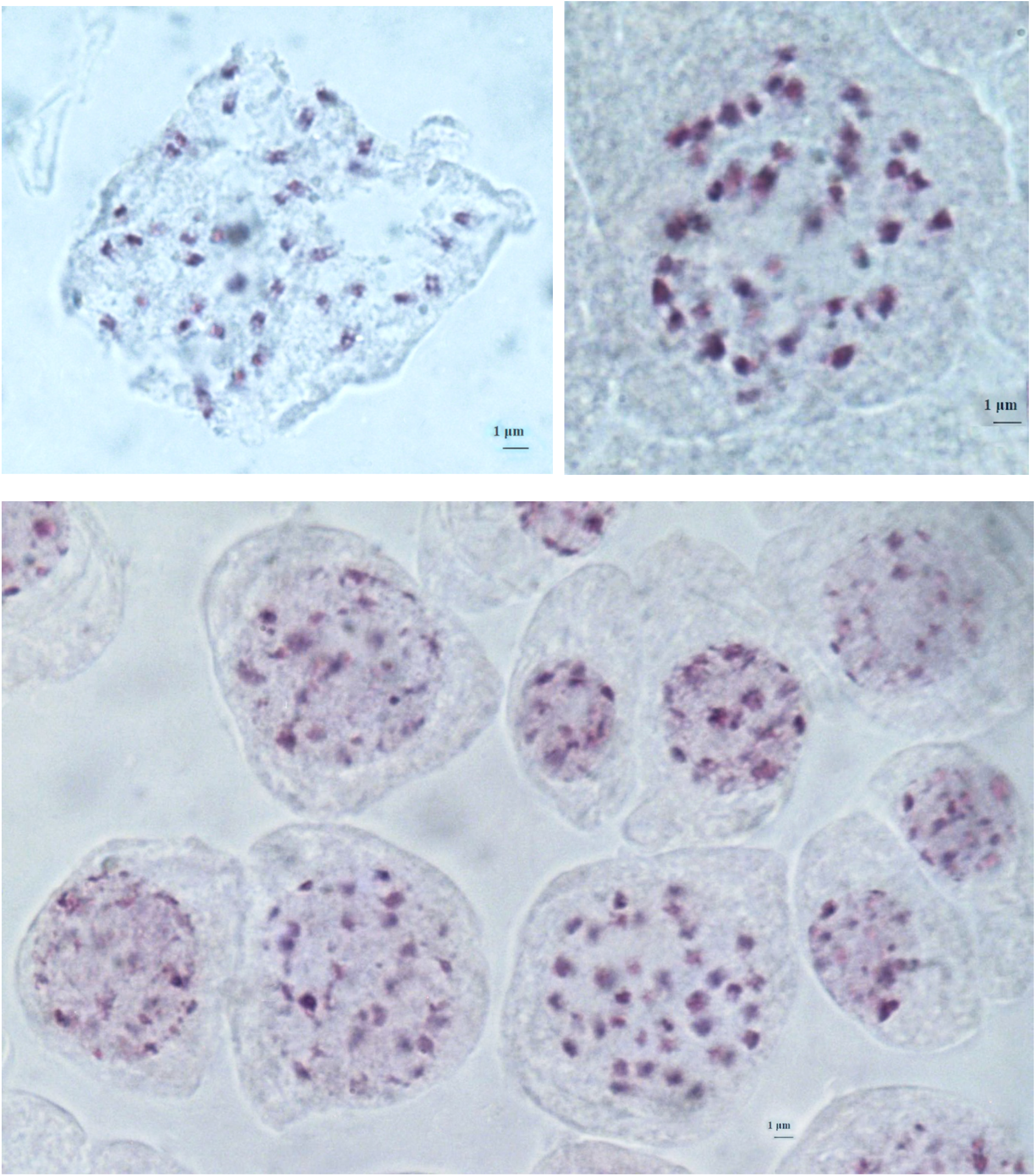
Chromosomes at somatic metaphase (Feulgen staining) of *× Pseudorhiza nieschalkii* nothosubsp. *siculorum* hybrid. 2n = 40 (±2), no. 28-2; no. 30-1; no. 31-4. Scale bar = 1 μm.

Sadly, an attempt made by Nora Anghelescu and Hajnalka Kertész five weeks later for more careful flower and pollination studies (in July 2020) were impeded by the complete destruction of the entire ‘protected’ area by uncontrolled cattle grazing. The hybrid was still relocated since natural landmarks and digital photos were taken in order to precisely mark the exact placement of the hybrid within the heathland. In autumn, millimetric explants (adventitious root tips) for chromosomal counts were collected in order to fully and precisely elucidate the parental origin of the hybrid.

Helped by the supportive morphometric results, we regard the karyotypical results as reliably demonstrating that the putative hybrid is indeed the F_1_ natural cross between *D. fuchsii* subsp. *sooana* and *P. albida* subsp. *tricuspis*. The status of this intergeneric hybrid as new to science was a further motivation to apply additional chromosomal analyses (as opposed to visual guesswork) in the hope of achieving a final and conclusive identification.

### Intergeneric hybridisation and speciation discussion

Over millions of years of evolution, hybridization had a major role in shaping the history of life on earth. The evolutionary history of a population is reflected in the genetic variation of its genomes. In natural populations, hybridization can act as an evolutionary engine by overcoming the reproductive barriers between populations.

Hybridisation is the crossing of two genotypically different parents, parent generations P_1_ × P_2_. The genes from P_1_ & P_2_ combine in the first resulted generation, named F_1_ (Soltis & Soltis, 2009). They will be present in the hybrid genotype and can be dominant, recessive or intermediate (Ramsey & Schemske, 1998). A new hybrid lineage is formed through parental genome mixing. The totality of all successful hybrid types that originate of the crossing of two parental taxa (natural species, not of hybrids) is called a nothotaxon. Nothotaxons may further progress as nothogenus, nothospecies or nothosubspecies. Hybridization is widespread, but the generation of a unique, natural hybrid lineage to occur is likely very rare. New hybrid lineages must establish reproductive isolation and a unique ecological niche in order to overcome genetic mixing and competition from parental species (Mayr, 1942). As a result, hybridisation was shown to have a significant role in speciation, generating new species with better genetic, adaptive variation (Arduino *et al.*, 1996).

### Contrasting pollination syndromes discussion

All species belonging to *Dactylorhiza* genus are nectar-deceit orchids. Their spurs are completely devoid of nectar and consequently, these *non-rewarding* species with *nectarless flowers do not offer any recompense* to the pollinating insects. Hence, their pollination is exclusively based on *deceit* and *mimicry*. All food-deceptive orchids exploit the pre-existing plant-pollinator relationships, especially the food foraging behaviour and achieve their pollination by deception. They are known as food-deceptive orchids and this particular pollination mechanism is classified as generalized food deception mechanism (Jersáková *et al.*, 2006). Despite the fact that they are rewardless, all *Dactylorhiza* species are *allogamous* (they do not self- or auto-pollinate) and depend entirely on insects for their *cross-pollination* and *seed production*. They are also generalist pollinators, usually pollinated by bees, bumble-bees, beetles, butterflies, flies, and share most of their pollinators with all the neighbouring nectariferous, rewarding plant species (reviewed in Claessens & Kleynen. 2011). In order to attract insects and successfully accomplish their pollination, they usually grow among rewarding plant species, which they very often mimic in one or more floral traits such as, inflorescence shape, flower colour, floral scents (that mimic the presence of nectar), nectar guides, spurs and pollen-like papillae. Little (1983) termed this type as ‘*mimicry based on naïveté*’.

Although most authors have argued that *Dactylorhiza* in general, offer no reward to the visitor, Dafni & Woodell (1986) demonstrated that some insects refuse to leave the nectarless flowers empty handed and, instead they start chewing the sugar-rich stigmatic exudates. *Dactylorhiza fuchsii*, the most intensively studied species for pollinators has been recorded as receiving visits from 35 insect species (Claessens & Kleynen, 2011; Dafni & Woodell, 1986; Bateman, 2017). Having taken in consideration that the heathland was populated by numerous Hymenopterans and Dipterans, it is very likely that the dominant pollinators of *D. fuchsii* subsp. *sooana* are diurnal various species of bumble-bee (Bombus genus), bees (Apis genus) and various butterflies.

*Pseudorchis* on the other hand, is a rewarding genus, which attracts its pollinators with its sweet, vanilla fragrance (secreted both during the day and the night) and recompenses them with large amounts of nectar. This pollination mechanism is known as generalised food foraging behaviour mechanism (Galizia *et al.*, 2005). As a result, *P. albida* subsp. *tricuspis* is able to attract a wide variety of diurnal pollinators such as bees, butterflies and beetles, especially during the hot summer days when the alpine plains are crowded with insects, all hungry and in search for food. Notably, the scent emission increases in the evening, thus attracting various crepuscular and nocturnal moths. The fruit set and seed production are very high in case of *P. albida* subsp. *tricuspis*, often over 90% (Detto, 1905; Ziegenspeck, 1936), which raises the possibility of spontaneous autogamy (Summerhayes, 1968; Nilsson, 1992; van der Cingel. 1995), particularly in cold alpine areas where, rather often, the pollinators are scarce (Jersáková *et al.*, 2011).

Within the hybrid’s neighbourhood, the parental species occurred in very close proximity of each other, approximately at a distance measuring 2-10 metres. This implies that the pollinating insects required a minimum travel distance between the parents, in order to successfully transport pollinaria (singular: pollinarium) between individuals and generate the hybrid. Since all three taxons display a considerable synchronicity in their flowering time, and, at least partially, may share the pollinator community, frequent exchange of pollen between the parental species was/is very likely.

Thus, the resulting hybrid proofs that even highly contrasting pollination syndromes such as generalized food deception and generalised food foraging behaviour mechanisms, are insufficient to stop the gene flow between two different orchid genera.

Since the resulted hybrid had significantly larger labellum than, but very similar to *P. albida* subsp. *tricuspis* parent, we speculate that prospective pollinating insects (shred from the parental pollinator communities) will perceive a flatter and proportionally wider landing stage in the hybrid than in the respective parent. Rather often, these character shifts are capable of modifying pollinator specificity, indicating a potential evolutionary future for the hybrid. Further data will explore whether the inheritance of such specific ‘character suites’ in intergeneric hybrids, impair, neutralise or enhance the functionality of these particular novel combinations of character states (Kretzschmar *et al.*, 2007).

In the same time, further studies will be needed to elucidate whether the hybrid is completely rewardless, similar to *D. fuchsii* subsp. *sooana* parent, or whether it is, at least partially, rewarding and secrets nectar in its, eventually, sweet vanilla-scented *‘tricuspis*-like’ flowers.

### Mycorrhizal associations discussion

The successful germination of at least one F_1_ seed and its development to maturity also merit special consideration, since most of the species belonging to *Dactylorhiza* and *Pseudorchis* genera (parental genera) require a period of approximately 4 years from seed germination to first appearance above ground (Fuchs & Ziegenspeck,1925; Rasmussen, 1995, 2003).

Both *D. fuchsii* (Bailarote *et al.*, 2012) and *P. albida* are widely viewed as mycorrhizal generalists, able to form partnerships with many fungi, thus having a much lower dependency than obligate mycoheterotrophs (Bateman *et al.*, 2017). *P. albida* is generally associated with a wide variety of fungi members of a polyphyletic basidiomycete group collectively called Rhizoctonia (Downie, 1959), which are typically found in photosynthetic orchids of open habitats (Rasmussen, 2002). Molecular research, based on cladistic analysis of DNA sequences, places Rhizoctonia within the family Ceratobasidiaceae (Moncalvo J-M *et al.*, 2006). Most of *Dactylorhiza* species are associated with strains of Ceratobasidium and Thanatephorus (anamorph Rhizoctonia) which have been isolated from the roots of adult individuals (Rasmussen, 1995, 2003).

Since the parental species share the same mycorrhizal partners/community, it is very likely that, should they be produced, the seeds of the hybrid might fall in close parental proximity and be stimulated to germinate by the same mycorrhizal fungi. It is well-known that the seeds (in this case the hybrid seeds) fall within the close vicinity of the parental plants and, by making use of the mycorrhizal fungi available, they often successfully germinate. The discussion of potential production of seed was made on the assumption that since the hybrid is a diploid (alloploid) individual (2n=40), is very likely to produce seeds by geitonogamous pollination (self-pollination).

It was very disappointing we were unable to complete the morphological study of the flowers, fruit and seeds, as the whole area was destroyed by cattle, a few weeks after the initial identification of the hybrid was done. The violent cattle intervention resulted in the grazing of the upper half all plants in the area. We managed to relocate the hybrid and identified its remains, namely the basal leaves and the lower half of the stem, which escaped by miracle, the hungry, voracious mouths of the herbivores.

### Maternity-Paternity Testing

Future research should imply extensive molecular analyses to establish the true maternal and parental origins/species of this intergeneric hybrid, i.e. (maternal parent) *D. fuchsii* subsp. *sooana*^♀^ × *P. albida* subsp. *tricuspis*^♂^ or (paternal parent) *D. fuchsii* subsp. *sooana*^♂^ × *P. albida* subsp. *tricuspis*^♀^).

These morphological and molecular investigations might also reveal the amount and direction of gene flow in orchids, which consequently generate specific enhanced characters in the hybrids relative the parental species. As mentioned, further field investigations will be needed to validate the fertility of this hybrid (even if such F_1_ plants are characterised by significant reduced fertility (Bateman *et al.*, 2017) and the presumed potentiality of its seeds to germinate and continue to produce new individuals beyond this first generation, F_1_ which, ultimately, might lead to the establishment of a new nothopopulation. In the same time, since the chromosomal numbers of the hybris and its parental species are very close (2n=40, 42) it would be interesting to show whether this F_1_ hybrid will be able to back-cross with either of its parental species and generate further new phenotypically distinct offspring.

### Could this unique hybrid be regarded as a new nothospecies?

According to strict etiquette it became common among botanists to officially describe all newly found hybrids as nothospecies or nothosubspecies, even though most are ephemeral (and, in most cases, infertile) and thus, a very small fraction of them eventually survive to develop stable nothospecies/nothopopulations (homoploid species) (Bateman *et al.*, 2017).

Despite these observations, we eventually chose to formally describe this hybrid between *D. fuchsii* subsp. *Sooana* and *P. albida* subsp. *tricuspis*, an *intergeneric hybrid, new to science*.

### × *Pseudorhiza nieschalkii* (Senghas) P.F.Hunt nothosubsp. *siculorum* H.Kertész & N.Anghelescu, 2020, nothosp. nov

#### Hybrid formula

*Dactylorhiza fuchsii* (Druce) Soó subsp. *sooana* (Borsos) Borsos **×***Pseudorchis albida* (L.) Á.Löve & D.Löve subsp. *tricuspis* (Beck) E. Klein

#### Diagnosis

Morphologically intermediary between the parents. Stem and leaves similar to *P. albida* subsp. *tricuspis* by the vividly green colour and the absence of spots on the leaves. The influence of *D. fuchsii* subsp. *sooana* parent may be seen in the whitish colour of the flowers and faint pinkish marks on the lateral sepals and the labellum. The marks are significantly fainter and resemble the scattered dots and lines found on *D. fuchsii* subsp. *sooana*’s labellum, which never form circular loops. The most distinctive feature is the three-lobed labellum, with all three lobes of equal length and width, most similar to *P. albida* subsp. *tricuspis* parent.

#### Description

Stem 28.5 centimetres, 3.2 millimetres in diameter, devoid of anthocyanins. Basal sheath present. Sheathing leaves 4, evenly distributed evenly on the stem, angled, laterally spreading, narrowly lanceolate, longest leaf 65 millimetres long, widest leaf 11 millimetres wide, longest placed basally/base of the stem. Leaves vivid green, unmarked, lacking pronounced apical hooding. Inflorescence 67 millimetres (23% of stem length), cylindrical, flat topped, fairly lax. Flowers 28. Labellum deeply three-lobed, width × length 5.1 × 4.2 millimetres to central lobe, 3.9 millimetres to lateral lobes. Inter-lobe sinuses deep, c.2.1-2.3 millimetres deep. Lateral lobes only slightly recurved, presenting partial margin crenulation. Labellar base colour yellow with a greenish tinge at spur base, while labellar surface became gradually pure white. Markings comparatively low-contrast pink dots and streaks, covering central region of labellum and the three lobes. Markings never form continuous loops. Labellar spur cylindrical, 3.4 millimetres long × 1.5 millimetres wide at mouth, 1.2 millimetres midway along length, slightly down-curved, shorter than the ovary. Lateral sepals lightly sprinkled with purple dots and streaks, spreading laterally and horizontally, relative to the loose hood formed by the median sepal and lateral petals. Basal bracts 16 millimetres, exceeding flowers, upper floral bracts 14 millimetres, both exceeding ovaries, top bracts lightly purple tinged with anthocyanin pigments.

#### Chromosome number

2n=40.

#### Flowering time

From mid-June to mid-July.

#### Locus classicus

*× Pseudorhiza nieschalkii* nothosubsp. *siculorum* is the only known plant, an F_1_ intergeneric hybrid, found within the Natura 2000 protected area ROSCI00090 Harghita – Mădăraş and ROSPA0034 Depresiunea şi Munţii Ciucului, in the peatbog area adjacent to Harghita Racoşului Peak, next to the mountain refuge, up to 1,700 meters altitude. GPS: 25.61451, 46.43298, currently in Harghita County Romania.

**Fig. 7.**
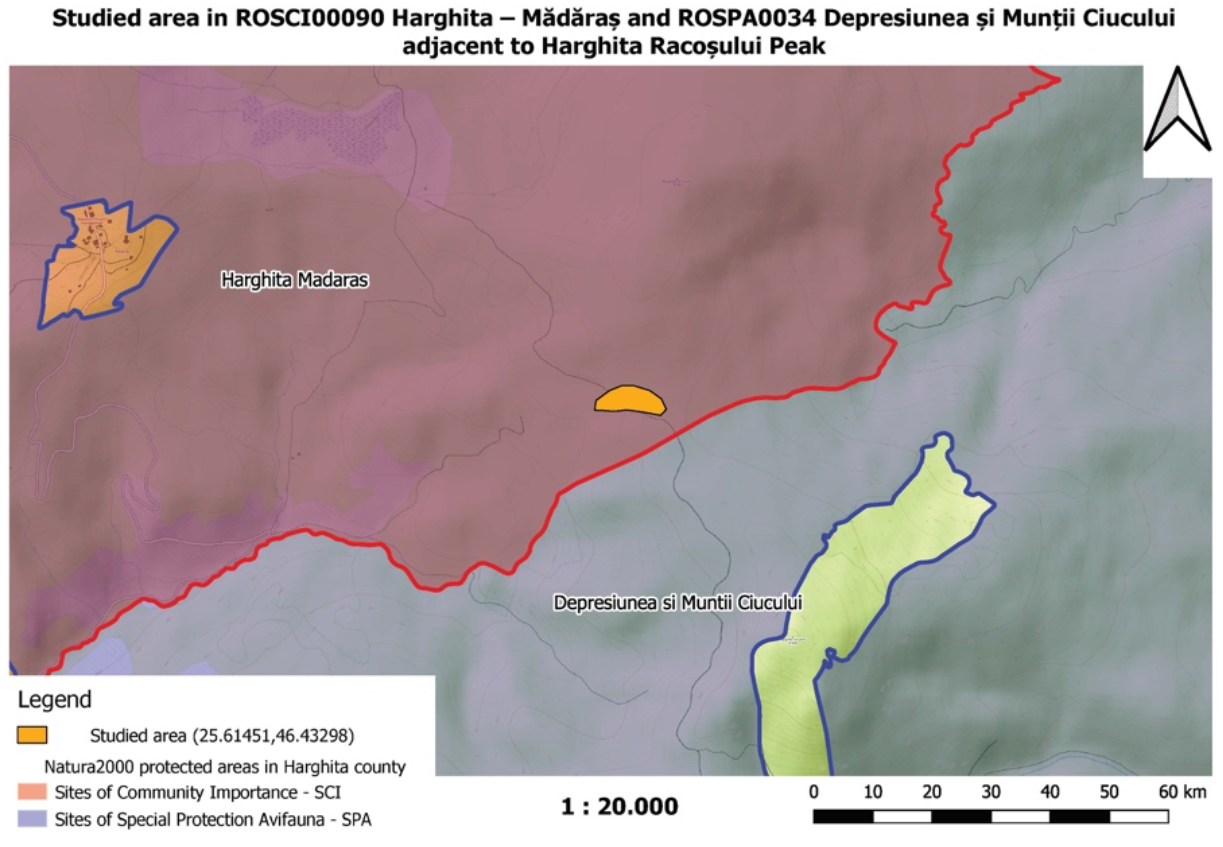
The distribution map of *× Pseudorhiza nieschalkii* nothosubsp. *siculorum* in Harghita – Mădăraş protected area that yields the holotype and only known plant (map used by permission of National Agency for Protected Areas Harghita Territorial Service)

#### Habitat

× *Pseudorhiza nieschalkii* nothosubsp. *siculorum* prefers a sunny, swampy meadow, next to mixed forest margin, on mildly acidic, nutrient poor substrate. The area under study was covered in moist substrate of *Sphagnum* L. bogs.

#### Holotype

The intergeneric hybrid *× Pseudorhiza nieschalkii* nothosubsp. *siculorum* represents the only individual ever reported, which was not herbalized. The holotypus was confined to printed digital images: 75 images taken by Hajnalka Kertész on the 30^th^ June (time: 08:37:19am), 3^rd^ of July and 30^th^ of June 2020 and 370 images taken by Nora Anghelescu on 3^rd^ of July and 30^th^ of June 2020, deposited in private image data bases. Photographs were taken but no voucher material was removed from so singular a plant.

#### Etymology

The intergeneric name, *Pseudorhiza*, is a compound term that originates from the fusion of the first and last parts of the scientific names of two distinct genera, *Pseudo*(rchis) and (Dactylo)*rhiza*. It refers to the rare cross between the species *Dactylorhiza fuchsii* subsp. *sooana* (Borsos) Borsos and *Pseudorchis albida* subsp. *tricuspis* (Beck) E.Klein, which were the only species found in the hybrid’s close proximity.

The nothospecies epithet, *nieschalkii*, was given in honor of the German botanist Albert Nieschalk (1904-1985), who discovered the hybrid *× Pseudorhiza nieschalkii* (*Dactylorhiza fuchsii* × *Pseudorchis albida*), in in 1947 in Germany at the ‘New Hagen’ near Niedersfeld in North Rhine-Westphalia (Eccarius, 2016), hence its potential vernacular name, Nieschalk’s Pseudorhiza.

The nothosubspecies epithet, *siculorum*, comes from the Latin name *Terra Siculorum* meaning *Siculorum County*, ad litteram meaning *of Siculorum*, a reference to the area where this unique orchid hybrid is endemic, which gave its vernacular name Pseudorhiza Siculorum. The nothosubspecies epithet, *siculorum*, was subsequently given to this new hybrid reflecting the identity of the region, Siculorum County, that yielded the holotype and only known plant. Terra Siculorum known as the Székely Land or Ţinutul Secuiesc is a historic and ethnographic area in Romania, inhabited mainly by Székelys. It includes the whole territories of Mureş, Harghita and Covasna counties and its main cultural centre is the city of Târgu Mureş, the largest settlement in the region.

## CONCLUSIONS

In order to elucidate the origin of the newly discovered hybrid, *× Pseudorhiza nieschalkii* nothosubsp. *siculorum*, we applied two different scientific methods, morphometrics and chromosomal counts plus one less scientific method, ‘expert intuition’, to unravel the history of the plant in question. As data accumulated, we eventually concluded with confidence that the plant is a spontaneous F_1_ hybrid generation that has *D. fuchsii* subsp. *sooana* and *P. albida* subsp. *tricuspis* as parents.

In the same time, we openly wonder how many reports of such hybrid occurrences received from various field workers or enthusiast botanists are in fact genuine hybrids and, contrarily, how many genuine hybrids pass unnoticed during field surveys because they resemble more closely one or other of their parents.

To conclude, we consider this novel combination an extremely fortunate find considering the exceptional geographic restriction/isolation of the parental species, in the fact that *D. fuchsii* subsp. *sooana*, exclusively confined to Harghita-Mădăraş protected area, favours the mildly acid soils (moist heathland), whereas *P. albida* subsp. *tricuspis* is typically a plant of soils that are moist to wet and neutral to alkaline (marshes, limestone grassland). Altogether, these contrasting requirements and reduced distribution of one parental species, offer very slim chances of the parental taxons to develop juxtaposed/sympatric populations. Harghita-Mădăraş natural reserve represents a unique exception.

## ACKNOWLEDGMENTS

We would like to thank Mr. Tőke Árpad & Mr. Demeter László, Director of the National Agency for Protected Areas Harghita Territorial Service for permission to investigate *Dactylorhiza* and *Pseudorchis* populations. We would also like to express our gratitude to orchidologist Prof. Helmut Presser, Germany for helpful discussion and information on species morphology, ecology and distribution, and to Prof. Paulina Anastasiu, Director of Dimitrie Brândză Botanical Garden, University of Bucharest, Romania for insightful advice. We are also grateful to the members of the Editorial Board for their suggestions and corrections. The work was privately funded by the authors.

## Notes

### Competing Interest Statement

The authors have declared no competing interest.

### Summary of Updates

Minor references and spelling errors were corrected.

## REFERENCES & BIBLIOGRAPHY

Arduino P, Verra F, Cianchi R, Rossi W, Corrias B & Bullini L. 1996. Genetic variation and natural hybridization between *Orchis laxiflora* and *Orchis palustris* (Orchidaceae). Pl. Syst. Evol. 202: 87. https://doi.org/10.1007/BF00985819.

Bailarote BC, Lievens B & Jacquemyn H. 2012. Does mycorrhizal specificity affect orchid decline and rarity? American Journal of Botany 99: 1655–1665.

Bateman R & Hollingsworth P. 2004. Morphological and Molecular Investigation of the Parentage and Maternity of *Anacamptis × albuferensis* (*A. fragrans* × *A. robusta*), a New Hybrid Orchid from Mallorca, Spain. Taxon 53 (1): 43–54. doi:10.2307/4135487.

Bateman RM & Denholm I. 1985. A reappraisal of the British and Irish dactylorchids, 1. The tetraploid marsh-orchids. Watsonia 14: 347–376.

Bateman RM & Denholm I. 2012. Taxonomic reassessment of the British and Irish tetraploid marsh-orchids. New Journal of Botany 2: 37–55. http://dx.doi.org/10.1179/2042349712Y.0000000004.

Bateman RM. 2001. Evolution and classification of European orchids: insights from molecular and morphological characters. Journal Europäischer Orchideen 33: 33–119.

Bateman RM. 2006a. Developing reliable tests for putative hybrids of bizarre parentage: a role for the BSBI? BSBI News 101: 17–20.

Bateman RM., Murphy ARM & Tattersall BG. 2017. × *Dactylodenia lacerta* (Orchidaceae): a morphologically cryptic hybrid orchid new to science from the Lizard Peninsula, Cornwall, New Journal of Botany, 7:2-3, 64–77. DOI: 10.1080/20423489.2017.1408189.

Borsos O. 1959. *D. fuchsii* subsp. sooana. Acta Bot. Acad. Sci. Hung. 5: 324

Borsos O. 1961. *D. fuchsii* subsp. *sooana.* First published in *Ann. Univ. Sci.* Budapest. Rolando Eötvös, Sect. Biol. 4: 76

Cavruc V. 2000. Archaeological Repertory of Harghita County, ISBN 973-99270-2-5.

Ciocârlan V. 2000: Flora ilustrată a României. Pteridophyta et Spermatophyta. Editura Ceres Bucureşti. 1138.

Ciocârlan V. 2009: Flora ilustrată a României. Editura Ceres Bucureşti. 1141 pag. ISBN 978-973-40-0817-9.

Claessens J & Kleynen J. 2011. Theflower of the European orchid: form and function. Voerendaal, The Netherlands: Published by the authors.

Council Directive 92/43/EEC of 21. Consolidated version: 01/07/2013.

Coyne JA & Orr HA. 2004. Speciation. Sunderland: Sinauer Associates.

Dafni A & Woodell SRJ. 1986. Stigmatic exudate and the pollination of *Dactylorhiza fuchsii* (Druce) Soó. Flora 178: 343–350.

Danihelka J, Chrtek J & Kaplan Z. 2012. Checklist of vascular plants of the Czech Republic Preslia. Casopsi Ceské Botanické Spolecnosti 84: 647–811.

De Angelli N & Anghelescu D. 2020. Orchids of Romania. Snagov, Romania. Published by the authors

Delforge P. 2006. Orchids of Europe, North Africa and the Middle East. A & C Black. ISBN-13: 978-0-7136-7525-2.

Detto, C. 1905. Blütenbiologische Untersuchungen I. Flora 94: 287–329.

Dev.adworks.ro. 2014. Natura 2000 - Harghita Mădăraş - Site of community importance.

Dihoru G & Negrean G. 2009: Cartea roşie a plantelor vasculare din România. Editura Academiei Române, Bucureşti, Romania.

Djordjević V, Tsiftsis S, Lakušić D, Jovanović S & Stevanović V. 2016: Factors affecting the distribution and abundance of orchids in grasslands and herbaceous wetlands. – Systematics and Biodiversity 14: 355–370.

Eunis.eea.europa.eu - Mădăraş Harghita (flora and fauna).

Fuchs A & Ziegenspeck H. 1925. Bau und Form der Wurzel der einheimi-scher Orchideen in Hinblick auf ihre Aufgaben. Botanisches Archiv 12: 290–379.

Govaerts, RHA. 2011. World checklist of selected plant families published update Facilitated by the Trustees of the Royal Botanic Gardens, Kew. [Cited as *Dactylorhiza fuchsii* subsp. *Sooana*.]

Hunt, PF. 1971. In: Orchid Rev. 79: 142.

Ielencz, M. 2005. Geography of Romania. University Publishing House. Bucharest, Romania.

Iucnredlist.org. 2020. The IUCN Red List of Threatened Species.

Jeřabková K. 2006. Ecological demands and optimal management of Pseudorchis albida. Master thesis in Czech, Czech University of Life Sciences, Prague.

Jersáková J & Malinová T. 2007. Spatial aspects of seed dispersal and seedling recruitment in orchids. New Phytologist 176: 237–241.

Jersáková J, Johnson SD & Kindlmann P. 2006. Mechanisms and evolution of deceptive pollination in orchids. Biol. Rev. 81: 219–235. doi:10.1017/S1464793105006986.

Jersáková J, Malinová T, Jeřabková K & Dötterl S. 2011: Biological Flora of the British Isles: Pseudorchis albida

(L.) Á. & D. Löve. – Journal of Ecology 99: 1282–1298.

Klein, E. 2000. *Pseudorchis albida* subsp. *tricuspis* (Beck) Klein stat. nov., eine weitgehend ü bersehene, calcicole, alpisch-boreale Sippe (Orchidaceae –Orchideae). Phyton 40: 141–159.

Kretzschmar H, Eccarius W & Dietrich H. 2007. The Orchid Genera Anacamptis, Orchis and Neotinea.

Kreutz, CAJ. 2014. Neue Erkentnisse zu den Orchideen Rumäniens. Ber. Arbeitskrs. Heim. Orchid. 31(2): 82–146.

Little, RJ. 1983. A review of floral food deception mimicries with comments on floral mutualism. In Handbook of Experimental Pollination Biology (eds. C. E. Jones and R. J. Little): 294–309. S & E Scientific and Academic Editions, New York.

Loya, V. 2015. *Dactylorhiza fuchsii* (Druce) Soó subsp. *sooana* (Borsos) Borsos in Transcarpathian flora. Sci. Bull. Uzhgorod Univ. (Ser. Biol.) Vol. 38-39.

Marcu O, Racz Z & Cioacă A. 1986. Harghita Mountains - tourist guide, Sport Tourism Publishing House, Bucharest, Romania.

Mayr, E. 1942. Systematics and the origin of species. New York: Columbia University Press.

Mikfalvi Z & Vifkori L. 1979. Harghita County, Monograph, Sport Turism Publishing House, Bucharest.

Moncalvo J-M et al. 2006. The cantharelloid clade: dealing with incongruent gene trees and phylogenetic reconstruction methods. Mycologia. 98 (6): 937–948. doi:10.3852/mycologia.98.6.937.PMID 17486970.

Moore DM. 1980: *Pseudorchis* SÉGUIER. In: Tutin T. G., Heywood V. H., Burges N. A., Moore D. M., Valentine D. H., Walters S. M. & Webb D. A. (eds.), Flora Europaea. Volume 5: Alismataceae to Orchidaceae (Monocotyledones). – Cambridge University Press, Cambridge, 332 pag.

Natura2000.mmediu.ro - Harghita Mădăraş Site of community importance.

Nilsson Ö. 1992. Pseudorchis albida ssp. *albida* - vityxne. Floravård I Jordbrukslandskapet Skyddsvärda (eds Ingelög T, Thor G, Hallingbäck T, Andersson R & Aronsson M): 266–267, SBT-förlaget, Lund, Sweden.

Oprea A. 2005: Lista critică a plantelor vasculare din România. Editura Universităţii Al. I. Cuza Iaşi, 668 pag.

Order of the Ministry of Environment and Sustainable Development No. 1964 of December 13, 2007, published in the Official Gazette of Romania, Part I, no. 98 and 98 bis of 7 February 2008.

Paucă A & Ştefureac Tr. 1972: *Leucorchis* E. MEYER. – In: SăVulescu T. (ed.), Flora Republicii Socialiste România 12, Editura Academiei Republicii Socialiste România, Bucureşti: 718–720.

Pedersen HÆ & Hedrén M. 2010. On the distinction of *Dactylorhiza baltica* and *D. pardalina* (Orchidaceae) and the systematic affinities of geographically intermediate populations. Nordic Journal of Botany 28: 1–12. http://dx.doi.org/10.1111/j.1756-1051.2009.00450.x.

Pisota I & D Bugă. 1976. Harghita County. Academy Publishing House. Bucharest, Romania.

Presser, H. 2002. Orchideen. Die Orchideen Mitteleuropas und der Alpen. Nikol. ISBN-13: 978-3933203540

Protectedplanet.net - Harghita Mădăraş Site of Community Importance (Habitats Directive).

Ramsey J & Schemske DW. 1998. Pathways, mechanisms, and rates of polyploid formation in flowering plants. Annu Rev Ecol Syst 29: 467–501.

Rasmussen HN & Whigham D. 1993. Seed ecology of dust seeds *in situ*: a new study technique and its application în terrestrial orchids. American Journal of Botany 80: 1374–1378.

Rasmussen, HN. 1995. Terrestrial orchids - from seed to mycotrophic plant. Cambridge University Press. ISBN 0521451655. Transfered to digital printing 2003.

Roskov Y, Kunze T, Orrell T, Abucay L, Paglinawan L, Culham A, Bailly N, Kirk P, Bourgoin T, Baillargeon G, Decock W, De Wever A & Didžiulis V. 2014. Species 2000 & ITIS Catalog of Life: 2014 Annual Checklist. Species 2000: Reading, UK.

Sârbu I, Ştefan N & Oprea A. 2013: Plante vasculare din România: determinator ilustrat de teren. – Editura Victor, Bucureşti, 1320 pp.

Soltis PS & Soltis DE. 2009. The role of hybridization in plant speciation. Annu Rev Plant Biol 60: 561–588

Stevens PF. 2001 onwards. Angiosperm Phylogeny Website. Version 14, July 2017 [Page last updated: 06/22/2019]; http://www.mobot.org/MOBOT/research/APweb/

Summerhayes VS. 1951, 1968. *Wild Orchids of Britain*. Collins, London, UK. The Board of Trustees of the Royal Botanic Gardens, Kew.

The International Plant Names Index and World Checklist of Selected Plant Families. 2020. Published on the Internet at http://www.ipni.org and http://apps.kew.org/wcsp/ © Copyright 2017 World Checklist of Selected Plant Families. http://creativecommons.org/licenses/by/3.0.

The Plant List. 2019. Version 1.1. Published on the Internet; http://www.theplantlist.org.

Van der Cingel NA. 1995. An Atlas of Orchid Pollination. European Orchids. A. A. Balkema, Rotterdam, Netherlands.

Ziegenspeck, H. 1936. Orchidaceae. Lebensgeschichte der Blütenpflanzen Mitteleuropas. Band 1, Abteilung 4. Eugen Ulmer, Stuttgart, Germany.

